# Accurately modeling biased random walks on weighted networks using *node2vec+*

**DOI:** 10.1101/2022.08.14.503926

**Authors:** Renming Liu, Matthew Hirn, Arjun Krishnan

## Abstract

**Motivation:** Accurately representing biological networks in a low-dimensional space, also known as network embedding, is a critical step in network-based machine learning and is carried out widely using *node2vec*, an unsupervised method based on biased random walks. However, while many networks, including functional gene interaction networks, are dense, weighted graphs, node2vec is fundamentally limited in its ability to use edge weights during the biased random walk generation process, thus under-using all the information in the network.

**Results:** Here, we present *node2vec+*, a natural extension of *node2vec* that accounts for edge weights when calculating walk biases and reduces to *node2vec* in the cases of unweighted graphs or unbiased walks. Using two synthetic datasets, we empirically show that *node2vec+* is more robust to additive noise than *node2vec* in weighted graphs. Then, using genome-scale functional gene networks to solve a wide range of gene function and disease prediction tasks, we demonstrate the superior performance of *node2vec+* over *node2vec* in the case of weighted graphs. Notably, due to the limited amount of training data in the gene classification tasks, graph neural networks such as GCN and GraphSAGE are outperformed by both *node2vec* and *node2vec+*

**Code Availability:** https://github.com/krishnanlab/node2vecplus_benchmarks

**Supplementary information:** Supplementary data are available at *Bioinformatics* online.

## 1 Introduction

Graphs and networks naturally appear in many real-world datasets including social networks and biological networks. The graph structure provides insightful information about the role of each node in the graph, such as protein function in a protein-protein interaction network (Liu *et al*., 2020; Krishnan *et al*., 2016). To more efficiently and effectively mine information from large-scale graphs with thousands or millions of nodes, several node embeddings methods have been developed (Cui *et al*., 2018; Hamilton *et al*., 2017). Among them, *node2vec* has been the top choice in bioinformatics due to its superior performance compared to many other methods (Ata *et al*., 2021; Yue *et al*., 2019). However, many biological networks, such as Greene *et al*. (2015); Johnson and Krishnan (2022), are dense and weighted by construction, which we demonstrate to be undesirable conditions for *node2vec* that can lead to sub-optimal performance.

*Node2vec* (Grover and Leskovec, 2016) is a second-order random walk based embedding method. It is widely used for unsupervised node embedding for various tasks, particularly in computational biology (Nelson *et al*., 2019), such as for gene function prediction (Liu *et al*., 2020), disease gene prediction (Peng *et al*., 2019; Ata *et al*., 2018), and essential protein prediction (Wang *et al*., 2021a; Zeng *et al*., 2021).Some recent works built on top of *node2vec* aim to adapt *node2vec* to more specific types of networks (Wang *et al*., 2021b; Valentini *et al*., 2021), generalize *node2vec* to higher dimensions (Hacker, 2021), augment *node2vec* with additional downstream processing (Chattopadhyay and Ganguly, 2020; Hu *et al*., 2020), or to study *node2vec* theoretically (Grohe, 2020; Davison and Austern, 2021; Qiu *et al*., 2018). Nevertheless, none of these follow-up works account for the fact that *node2vec* is less effective for weighted graphs, where the edge weights reflect the (potentially noisy) similarities between pairs of nodes. This failing is due to the inability of *node2vec* to differentiate between small and large edges connecting the previous vertex with a potential next vertex in the random walk, which subsequently causes less accurate modeling of the intended walk bias.

Meanwhile, another line of recent works on graph neural networks (GNNs) have shown remarkable performance in prediction tasks that involve graph structure, including node classification (Bronstein *et al*., 2021; Zhang *et al*., 2021; Wu *et al*., 2021). Although GNNs and embedding methods like *node2vec* are related in that they both aim at projecting nodes in the graph to a feature space, two main differences set them apart. First, GNNs typically require labeled data while embedding methods do not. This label dependency makes the embeddings generated by a GNN tied to the quality of the labels, which in some cases, like in biological networks, are noisy and scarce. Second, GNNs typically require node features as input to train, which are not always available. In the absence of given node features, one needs to generate them and often GNN algorithms in this case use trivial node features such as the constant features or node degree features. These two differences give node embedding methods a unique place in node classification, apart from the GNN methods.

Here, we propose an improved version of *node2vec* that is more effective for weighted graphs by taking into account the edge weight connecting the previous vertex and the potential next vertex. The proposed method *node2vec+* is a natural extension of *node2vec*; when the input graph is unweighted, the resulting embeddings of *node2vec+* and *node2vec* are equivalent in expectation. Moreover, when the bias parameters are set to neutral, *node2vec+* recovers a first-order random walk, just as *node2vec* does. Finally, we demonstrate the superior performance of *node2vec+* through extensive benchmarking on both synthetic datasets and network-based gene classification datasets using various functional gene interaction networks. *Node2vec+* is implemented as part of PecanPy (Liu and Krishnan, 2021) and is available on GitHub: https://github.com/krishnanlab/PecanPy.

## 2 Method

We start by briefly reviewing the *node2vec* method. Then we illustrate that *node2vec* is less effective for weighted graphs due to its inability to identify *out* edges. Finally, we present a natural extension of *node2vec* that resolves this issue.

### 2.1 *Node2vec* overview

In the setting of node embeddings, we are interested in finding a mapping *f* : *V* → ℝ^*d*^ that maps each node *v* ∈ *V* to a *d*-dimensional vector so that the mutual proximity between pairs of nodes in the graph is preserved. In particular, a random walk based approach aims to maximize the probability of reconstructing the neighborhoods for any node in the graph based on some sampling strategy *S*. Formally, given a graph *G* = (*V, E*) (the analysis generalizes to directed and/or weighted graphs), we want to maximize the log probability of reconstructing the sampled neighborhood 𝒩_*S*_(*u*) for each *u* ∈ *V* :

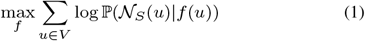

Under the conditional independence assumption, and the parameterization of the probabilities as the softmax normalized inner products (Grover and Leskovec, 2016; Mikolov *et al*., 2013b), the objective function above simplifies to:

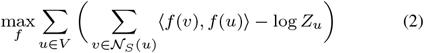

In practice, the partition function *Z*_*u*_ = ∑_*v*∈*V*_ ⟨*f* (*u*), *f* (*v*) ⟩ is approximated by negative sampling (Mikolov *et al*., 2013a) to save computational time. Given any sampling strategy *S*, equation (2) can find the corresponding embedding *f*, which is achieved in practice by feeding the random walks generated to the skipgram with negative sampling (Mikolov *et al*., 2013b).

*Node2vec* devises a second order random walk as the sampling strategy. Unlike a first order random walk (Perozzi *et al*., 2014), where the transition probability ℙ (*c*_*i*_|*c*_*i*−1_) depends only on the current vertex *c*_*i*−1_, a second order random walk depends also on the previous vertex *c*_*i*−2_, with transition probability ℙ (*c*_*i*_|*c*_*i*−1_, *c*_*i*−2_). It does so by applying a bias factor *α*_*pq*_(*c*_*i*−2_, *c*_*i*_) to the edge (*c*_*i*_, *c*_*i*−1_) ∈ *E* that connects the current vertex and a potential next vertex. This bias factor is a function that depends on the relation between the previous vertex and the potential next vertex, and is parameterized by the *return* parameter *p*, and the *in-out* parameter *q*. For the ease of notation, in the following, we denote *x* = *c*_*i*_ as the potential next vertex, *v* = *c*_*i*−1_ as the current vertex, and *t* = *c*_*i*−2_ as the previous vertex (see Figure 1). In this way, the random walk can be generated based on the following transition probabilities:

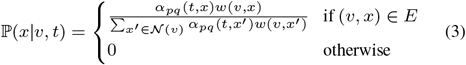

where the bias factor is defined as:

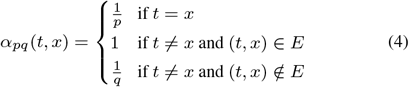

**Fig. 1.**
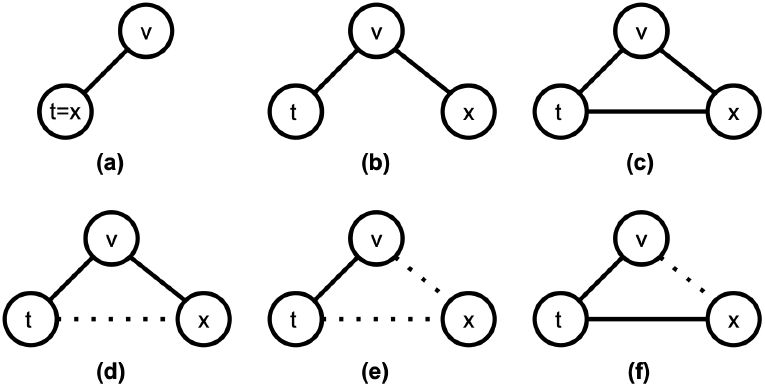
Illustration of different settings of return and in-out edges. The solid and dotted lines represent edges with large and small edge weights, respectively.

According to this bias factor, *node2vec* differentiates three types of edges: 1) the *return* edge, where the potential next vertex is the previous vertex (Figure 1a); 2) the *out* edge, where the potential next vertex is *not* connected to the previous vertex (Figure 1b); and 3) the *in* edge, where the potential next vertex is connected to the previous vertex (Figure 1c). Note that the first order (or unbiased) random walk can be seen as a special case of the second order random walk where both the *return* parameter and the *in-out* parameter are set to neutral (*p* = 1, *q* = 1).

We now turn our attention to weighted networks, where the edge weights are not necessarily zeros or ones. Consider the case where *t* and *x* are connected, but with a small weight (Figure 1d), i.e. (*t, x*) ∈ *E* and 0 < *w*(*t, x*) ≪ 1. According to the definition of the bias factor, no matter how small *w*(*t, x*) is, (*v, x*) would always be considered as an *in* edge. Since in this case *t* and *x* are barely connected, (*v, x*) should in fact be considered as an *out* edge. In the extreme case of a fully connected weighted graph, where (*u, v*) ∈ *E* for all *u, v* ∈ *V, node2vec* completely loses its ability to identify *out* edges.

Thus, *node2vec* is less effective for weighted networks due to its inability to identify potential *out* edges where the terminal vertex *x* is loosely connected to a previous vertex *t*. Next, we propose an extension of *node2vec* that resolves this issue, by taking into account of the edge weight *w*(*t, x*) in the bias factor.

### 2.2 Node2vec+

The main idea of extending *node2vec* is to identify potential out edges (*v, x*) ∈ *E* coming from node *t*, where *x* is loosely connected with *t*.Intuitively, we can determine the “looseness” of (*v, x*) based on some threshold edge value. However, given that the distribution of edge weights of any given node in the graph is not known *a priori*, it is hard to come up with a reasonable threshold value for all networks. Instead, we define the “looseness” of (*v, x*) based on the edge weight statistics for each node *v*.

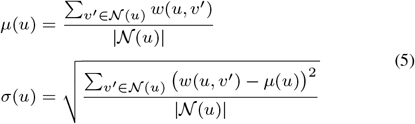

Formally, we first define *µ*(*v*) and *σ*(*v*) for *v* ∈ *V* to be the average and the standard deviation of the edge weights connecting *x*, respectively (Eqn 5). Then, we say *v* ∈ *V* is *γ-loosely connected* (or simply *loosely connected* if *γ* = 0) to *x* if 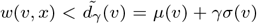, where *d*_*γ*_ (*v*) is the *noisy edge threshold* for node *v* parameterized by *γ* ∈ ℝ. Intuitively, we would like to treat an edge as being “not connected” if it is “small enough”. Note that in practice, because the graphs of interest only contain positive edge weights, we restrict the *noisy edge threshold* to be non-negative, i.e. 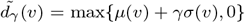 We also consider *v* being *γ-loosely connected* to *x* if (*v, x*) → *E*, as a slight abuse of notation. Finally, an (directed or undirected) edge (*v, x*) is *γ-loose* if *v* is *γ*-loosely connected to *x*, and otherwise it is *γ-tight*. Without loss of generality, we consider the case of *γ* = 0 in the subsequent sections to simplify the notion of *looseness*.

Based on the definition of looseness of edges, and assuming *t* ≠ *x*, there are four types of (*v, x*) edges (see Figure 1, (c-f)). Following *node2vec*, we categorize these edge types into *in* and *out* edges. Furthermore, to prevent amplification of noisy connections, we added one more edge type called the *noisy* edge, which is always suppressed.

#### 2.2.1 *Out* edge (1b, 1d)

As a direct generalization to *node2vec*, we consider (*v, x*) to be an *out* edge if (*v, x*) is tight and (*x, t*) is loose. The *in-out* parameter *q* then modifies the out edge to differentiate “inward” and “outward” nodes, and subsequently leads to Breadth First Search or Depth First Search like searching strategies (Grover and Leskovec, 2016). Unlike *node2vec*, however, we further parameterize the bias factor *α* based on 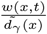. Any choice of monotonic function should work, but we choose to use the linear interpolation in this study for simplicity and leave it as future work to explore more sophisticated interpolation functions such as the sigmoidal functions. Specifically, for an *out* edge (*v, x*), the bias factor is computed as 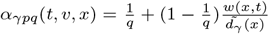. Thus the amount of modification to the *out* edge depends on the level of looseness of (*x, t*). When *w*(*x, t*) = 0, or equivalently (*x, t*) → *E*, the bias factor for (*v, x*) is 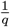, same as that defined in *node2vec*.

#### 2.2.2 *Noisy* edge (1e)

We consider (*v, x*) to be a *noisy* edge if both (*v, x*) and (*x, t*) are loose. Heuristically, the *noisy* edges are not very informative and thus should be suppressed regardless of the setting of *q* to prevent amplification of noise. Thus, the bias factor for a *noisy* edge is set to be min 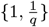.

#### 2.2.3 *In* edge (1c, 1f)

Finally, we consider (*v, x*) to be an *in* edge if (*x, t*) is tight, regardless of *w*(*v, x*). The corresponding bias factor is set to neutral as in *node2vec*.

Combining the above, the bias factor for *node2vec+* is defined as follows:

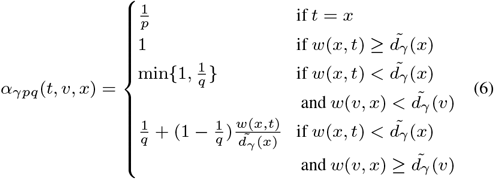

Note that the last case in equation (6) includes cases where (*x, t*) → *E*. Based on the biased random walk searching strategy using this bias factor, the embedding can be generated accordingly using (2). One can verify, by checking equation (6), that this is indeed a natural extension of *node2vec* in the sense that

- For an unweighted graph, the *node2vec+* is equivalent to *node2vec*.
- When *p* and *q* are set to 1, *node2vec+* recovers a first order random walk, same as *node2vec* does.

Finally, by design, *node2vec+* is able to identify potential *out* edges that would have been obliviated by *node2vec*.

## 3 Experiments

### 3.1 Synthetic datasets

We start by demonstrating the ability of *node2vec+* to identify potential out edges in weighted graphs using a barbell graph and the hierarchical cluster graphs. For simplicity, we fix *γ* = 0 for all experiments in this section.

#### 3.1.1 Barbell graph

A barbell graph, denoted as *B*, is constructed by connecting two complete graphs of size 20 with a common bridge node (Figure 2a). All edges in *B* are weighted 1. There are three types of nodes in *B*, 1) the bridge node; 2) the peripheral nodes that connect the two modules with the bridge node; 3) the interior nodes of the two modules. By changing the *in-out* parameter *q, node2vec* could put the peripheral nodes closer to the bridge node or interior nodes in the embedding space.

**Fig. 2.**
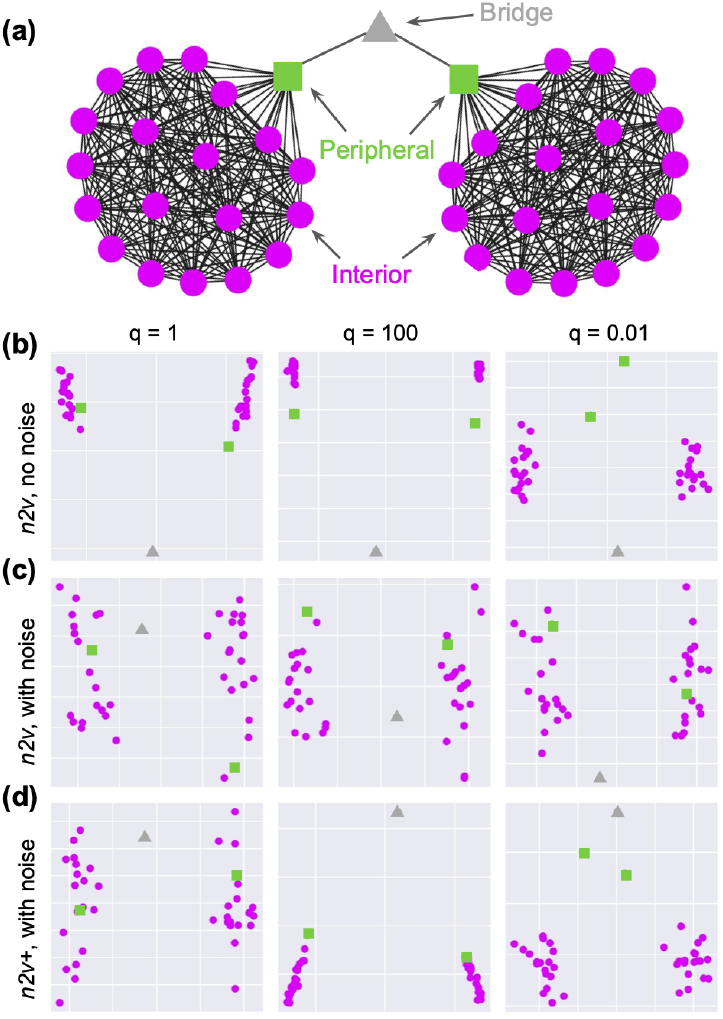
Barbell graph. (a) Illustration of the barbell graph, and three different types of nodes. (b)-(d) Embeddings of the barbell graph (or the noisy version of it), with three different settings of q = [1, 100, 0.01], and the color coding is the same as that in (a).

When *q* is large, *node2vec* suppresses the *out* edges, e.g., an edge connecting a peripheral node to the bridge node, coming from an interior node. Consequently, the biased random walks are restricted to the network modules. In this case, the transition from the peripheral nodes to the bridge node becomes less likely compared to a first-order random walk, thus pushing the embeddings between the bridge node and the peripheral nodes away from each other. Conversely, when *q* is small, the transition between the peripheral nodes and the bridge node is encouraged. In this case, the embeddings of the bridge node and the peripheral nodes are pulled together. To see this, we run *node2vec* with fixed *p* = 1, and three different settings of *q* = [1, 100, 0.01]. Indeed, for *q* = 100, *node2vec* tightly clusters interior nodes and pushed the bridge node away from the peripheral nodes, and for *q* = 0.01, the peripheral nodes are pushed away from the interior nodes (Figure 2b).

Next, we perturb the barbell graph by adding loose edges with edge weights of 0.1, making the graph fully connected. This perturbed barbell graph is denoted 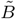. As expected, *node2vec* failed to make use of the *q* parameter (Figure 2c), since none of the edges are identified as an *out* edge. On the other hand, *node2vec+* can pick up potential *out* edges and thus qualitatively recovers the desired outcome (Figure 2d). Note that both *node2vec* and *node2vec+* have similar results for 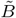 when *q* = 1. This confirms that *node2vec+* and *node2vec* are equivalent when *p* and *q* are set to neutral, corresponding to embedding with unbiased random walks. Finally, when using non-neutral settings of *q, node2vec+* is able to suppress some noisy edges, resulting in less scattered embeddings of the interior nodes (Figure 2d).

#### 3.1.2 Hierarchical CLUSTER graph

We use a modified version of the CLUSTER dataset (Dwivedi *et al*., 2020) to further demonstrate the advantage of the *node2vec+* due to identifying potential *out* edges. Specifically, the hierarchical cluster graph K3L2 contains *L* = 2 levels (3 including the root level) of clusters, and each parent cluster is associated with *K* = 3 children clusters (Figure 3a). There are 30 nodes in each cluster, resulting in a total of 390 nodes. To generate the hierarchical cluster graph, we first generate point clouds via a Gaussian process in a latent space so that the Euclidean distance between two points from two sibling clusters is about twice 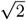 to be precise) the expected Euclidean distance from one of the two points to a point in the parent cluster, which is set to be 1. The noisiness of the clusters is controlled by the parameter *σ*, which is set to 0.1 by default. These data points are then turned into a fully connected weighted graph using a RBF kernel (see supplement). We consider two different tasks (Figure 3a), (1) *cluster classification*: identifying individual cluster identity of each node in the graph, and (2) *level classification*: identifying the level to which the clusters correspond to. We split the nodes into 10% training and 90% testing and use the multinomial logistic regression model with l2 regularization for prediction. The evaluation process, including the embedding generation, is repeated ten times, and the final results are reported by Macro F1 scores.

**Fig. 3.**
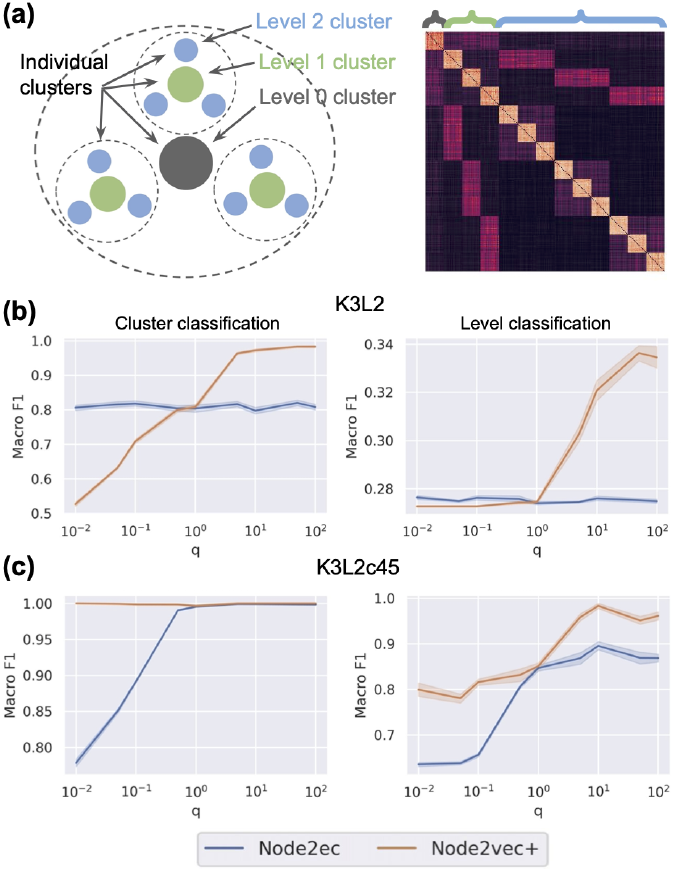
Hierarchical CLUSTER graph classification task. (a) Illustrations of the K3L2 hierarchical clusters. Left: top-down view of the clusters. Right: adjacency matrix of K3L2; colored brackets indicate the corresponding cluster levels of the nodes. (b) Classification evaluation on K3L2. (c) Classification evaluation on K3L2c45.

As shown in Figure 3b, the performance of *node2vec* is not affected by the *q* parameter because the graph is fully connected. Meanwhile, *node2vec+* achieves significantly better performance than *node2vec* for large *q* settings for both tasks, demonstrating the ability of *node2vec+* to identify potential *out* edges and use this information to perform localized biased random walks. Similar results are observed on a couple different hierarchical cluster graphs K3L3, K5L1, and K5L2 (see supplement).

On the other hand, one might suspect that the issue with fully connected graph can be alleviated by sparsifying the graph based on certain edge weight thresholds. Such an approach is widely adopted as a post processing step for constructing functional gene interaction networks. Here, we show that even after sparsifying the graph aggressively, *node2vec+* still outperforms *node2vec*. In particular, we sparsify the K3L2 graph using the edge weight threshold 0.45, which is the largest value that keeps the graph connected. We then perform the same evaluation analysis as above on this sparsified graph K3L2c45. In this case, *node2vec* indeed perform significantly better than before the sparsification for both tasks. Nonetheless, *node2vec+* achieves even better performance, still out-competing node2vec (Figure 3c).

Finally, we conduct a fine-grained evaluation analysis, showing that node2vec+ consistently outperforms node2vec under a wide range of conditions including edge threshold, train-test ratio, and noise-level (see supplement).

### 3.2 Real world datasets

Our main motivation of developing *node2vec+* stems from the fact that many functional gene interaction networks are dense and weighted. To systematically evaluate the ability of *node2vec+* to embed such biological networks, we consider various challenging gene classification tasks, including gene function and disease gene predictions. Furthermore, we devise experiments with previously benchmarked datasets BlogCatalog and Wikipedia (Grover and Leskovec, 2016) and confirm that *node2vec+* performs equal to or better than *node2vec* depending on whether the network is weighted (see supplement).

#### 3.2.1 Datasets

##### Human functional gene interaction networks

We consider functional gene interaction networks, which is a broader class of gene interaction networks that are routinely used to capture gene functional relationships.

- **STRING** (Szklarczyk *et al*., 2021) is an integrative gene interaction network that combines evidence of protein interactions from various sources, such as text-mining, high-throughput experiments, and etc.
- **HumanBase-global** is a tissue-naive version of the HumanBase (Greene *et al*., 2015) tissue-specific networks (previously known as GIANT), which are constructed by integrating hundreds of thousands of publicly available gene expression studies, protein-protein interactions, and protein-DNA interactions via a Bayesian approach, calibrated against high-quality known functional gene interactions.
- **HumanBaseTop-global** is a sparsified version of HumanBase-global that eliminates all edges below the prior of 0.1.

##### Multi-label gene classification tasks

We follow the procedure detailed in (Liu *et al*., 2020) to prepare the multi-label gene classification datasets. More specifically, we prepare two collections of gene classification tasks (each is called a gene set collection):

- **GOBP**: Gene function prediction tasks derived from the Biological Processes gene sets from the Gene Ontology (Consortium, 2018).
- **DisGeNet**: Disease gene prediction tasks derived from the disease gene sets from the DisGeNet database (Piñero *et al*., 2016).

After filtering and cleaning up the raw gene set collections, we end up with ∼45 functional gene prediction tasks and ∼100 disease gene prediction tasks (Table 1). These gene classification tasks are challenging primarily due to the scarcity of the labeled examples, with on average 100 and 200 positive examples per task for GOBP and DisGeNet, respectively, relative to the (order of) tens of thousands of nodes in the networks.

**Table 1.** Number of tasks (i.e., gene sets or node classes) for each combinations of network and gene set collection. The number in the parenthesis is the average number of positive examples.

|  | GOBP | DisGeNet |
| --- | --- | --- |
| HumanBase | 46 (98.3) | 103 (225.9) |
| STRING | 41 (100.0) | 97 (221.5) |

We split the genes into 60% training, 20% validation, and 20% testing according to the level at which they have been studied in the literature (based on the number of PubMed publications associated with each gene). In particular, the top 60% most well-studied genes are used for training; the 20% least-studied genes are used for testing, and the rest of the genes are used for validation. For GNNs, we report the test scores at the epoch where the best validation score is achieved.

#### 3.2.2 Baseline methods

We exclude several popular node embedding methods such as DeepWalk (Perozzi *et al*., 2014), LINE (Tang *et al*., 2015), and GraRep (Cao *et al*., 2015), from our main analysis, as it has been shown previously in various contexts (Grover and Leskovec, 2016; Ata *et al*., 2021; Yue *et al*., 2019) that *node2vec* is superior.

On the other hand, we include two popular GNNs, GCN (Kipf and Welling, 2016) and GraphSAGE (Hamilton *et al*., 2017) in our comparison. Both methods have shown exceptional performance on many node classification tasks, but their performance on the gene classification tasks here are still less well studied. For GraphSAGE, we consider the full-batch training strategy with mean pooling aggregation following the Open Graph Benchmark (Hu *et al*., 2021).

#### 3.2.3 Experiment setup

##### Evaluation metric

Following (Liu *et al*., 2020), we use the log_2_ 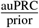 as our evaluation metric, which represents the log_2_ fold change of the average precision compared to the prior. This metric is more suitable than other commonly used metrics like AUROC as it corrects for the class imbalance issue that is prevalent in the gene classification tasks here, as well as emphasizes the correctness of top predictions.

##### Tuning embeddings parameters

For *node2vec* and node2vec+, we train a one vs rest logistic regression with l2 regularization using the embeddings learned. The parameters for embeddings including dimension, window-size, walk-length, and number of walks per node are set to 128, 10, 80, and 10, respectively, by default. We tune the hyperparameters for *node2vec* (*p, q*) and for *node2vec+* (*p, q, γ*) via grid search using the validation sets. To keep the grid search budget comparable, we search *p* and *q* over {0.01, 0.05, 0.1, 0.5, 1, 5, 10, 50, 100}^2^ for *node2vec* (*n* = 81); we search *p* and *q* over {0.01, 0.1, 1, 10, 100}^2^, together with *γ* ∈ {0, 1, 2} for *node2vec+* (*n* = 75).

##### Tuning GNN parameters

For both GNNs, we train one model for each combination of a network and a gene set collection in an end-to-end fashion. The architectures are fixed to five hidden layers with hidden dimension of 128. Since the gene interaction networks here do not come with node features, we use constant feature for GCN and degree feature for GraphSAGE, respectively. We use the Adam optimizer (Kingma and Ba, 2014) to train the GNNs with 100, 000 max number of epochs. The learning rates are tuned via grid search from 10^−5^ to 10^−1^ based on the validation performance. The optimal learning rates that result in decent convergence rate without diverging are 0.01 and 0.0005 for GCN and GraphSAGE, respectively (see supplement).

#### 3.2.4 Experimental results

##### Tuning *γ* significantly improves performance for dense graph

The *γ* parameter in *node2vec+* (see 2.2) controls the threshold of distinguishing *in* edges and and *out* edges. A small or negative valued *γ* considers most non-zero edges as *out* edges. Conversely, a large valued *γ* identifies less *out* edges. When the input graph is noisy and dense, assigning a larger *γ* (e.g., 1) can act as a stronger denoiser to suppress spurious *out* edges. Figure 4 compares the gene classification test performance between *γ* = 0 and *γ* = {1, 2} with optimally tuned *p, q* using the HumanBase-global network. Higher testing scores are achieved by larger *γ* settings, illustrating that, to properly “denoise” the fully connected weighted graph HumanBase-global, we need to increase the noisy edge thresholds. On the contrary, the difference in performance due to the *γ* settings is less pronounced for sparse networks like HumanBaseTop-global and STRING (see supplement).

**Fig. 4.**
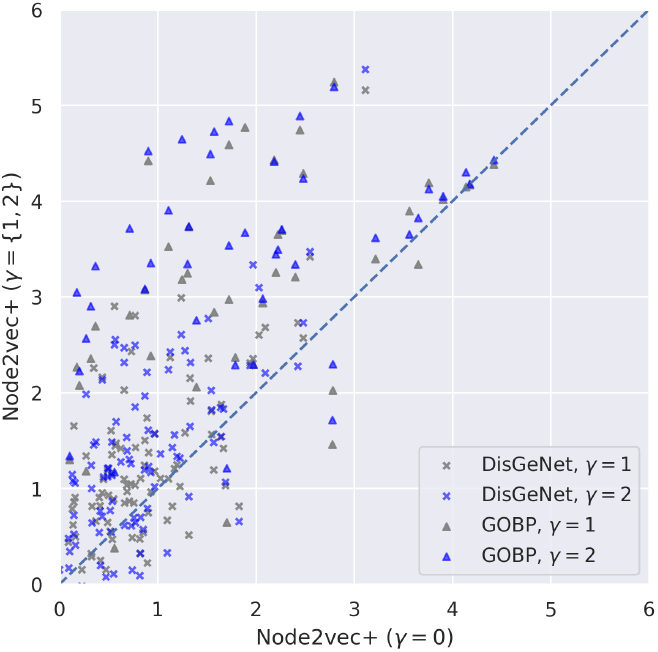
Comparison of different *γ* settings in node2vec+ using HumanBase-global. Each dot represents the testing performance 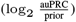 of a specific gene set, with optimally tuned *p* and *q* settings.

##### GNN methods performs worse than *node2vec(+)*

In all settings, *node2vec+* significantly outperforms both GNN methods (Figure 5). Particularly, for STRING network, both *node2vec* and *node2vec+* outperform the two GNNs by a large margin. The sub-optimal GNN performance here illustrates that, despite being powerful neural network architectures that can leverage the graph structures, GNNs by themselves cannot learn effectively given limited amount of labeled examples. On the contrary, the embedding processes of *node2vec(+)* are task agnostic and can be carried out effectively without labels. These results indicate that gene classification tasks based on gene interaction network are more effectively solved by unsupervised shallow embedding methods than GNNs.

**Fig. 5.**
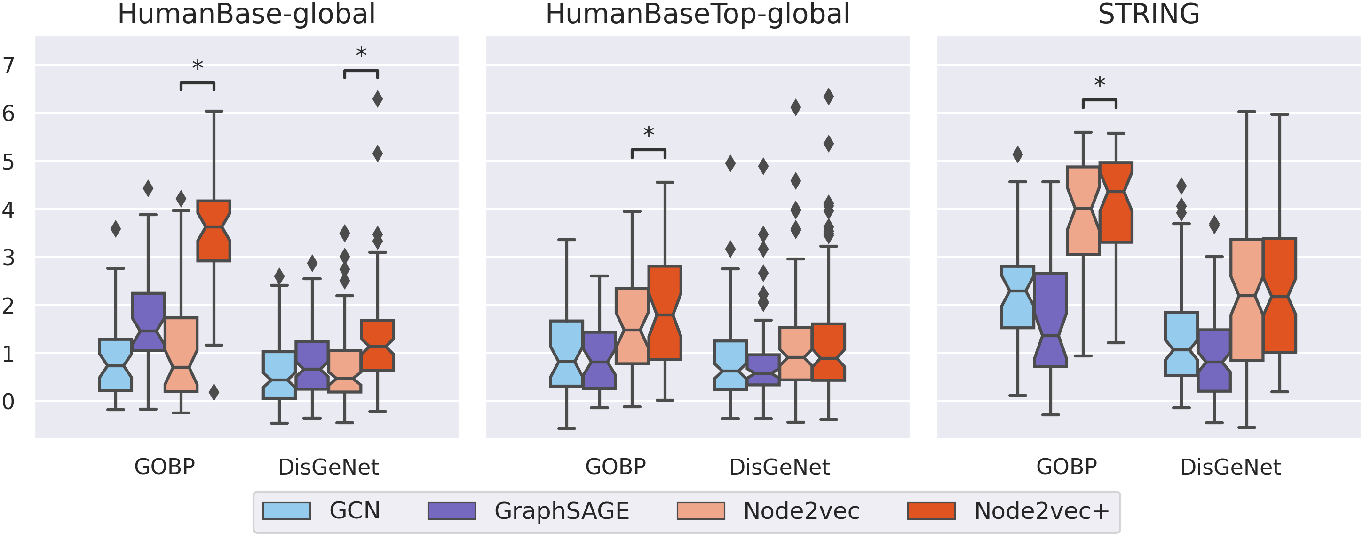
Gene classification tasks using protein-protein interaction networks. Each point in a boxplot represents the final test score for a specific task (gene set) in the gene set collection. Starred (*) pairs indicate that the performance between node2vec and node2vec+ are significantly different (Wilcoxon p-value *<* 0.05).

##### *nod2vec+* matches or outperforms *node2vec*

*node2vec+* significantly outperforms *node2vec* (Wilcoxon paired test (Wilcoxon, 1945) p-val < 0.05) except for the DisGeNet tasks using HumanBaseTop-global and STRING networks, in which cases the two methods perform equally (Figure 5). The performance differences are especially pronounced when using the fully connected and noisy HumanBase-global network, demonstrating *node2vec+*’s ability to learn robust node representations in the presence of noise. Nevertheless, when the network is less dense (e.g., HumanBaseTop-global), *node2vec+* is still able to perform at least as well as *node2vec*, indicating that *node2vec+* is overall a good replacement of *node2vec*.

#### 3.2.5 Tissue-specific functional gene classification

A key feature of functional gene interaction networks constructed using gene expression data is capturing biological context specificity, such as tissue-specificity provided by the HumanBase networks. Thus, we further demonstrate the use case of *node2vec+* using tissue-specific functional gene classification tasks derived from (Zitnik and Leskovec, 2017). After processing, there are 25 tissue-specific functional gene classification tasks, with 12 different tissues found in the HumanBase database. We follow a similar experimental setup as above and for each tissue-specific functional gene classification task we report the followings:(1) *matched*: the prediction performance using the corresponding tissue-specific network; (2) *other*: the average prediction performance using tissue specific networks other than the corresponding tissue; (3) *global*: the prediction performance using the tissue-naive network.

Figure 6 shows that *node2vec+* outperforms *node2vec* in most scenarios, especially when using the full HumanBase networks. In particular, *node2vec+*, using the *matched* tissue-specific full networks for the given functional gene classification tasks, results in significantly better performance than using *other* (unrelated) tissue-specific networks, as well as the *global* (tissue-naive) network. On the contrary, *node2vec* cannot fully utilize the tissue-specific networks, as indicated by the lack of difference in performance between *matched* and *global* networks.

**Fig. 6.**
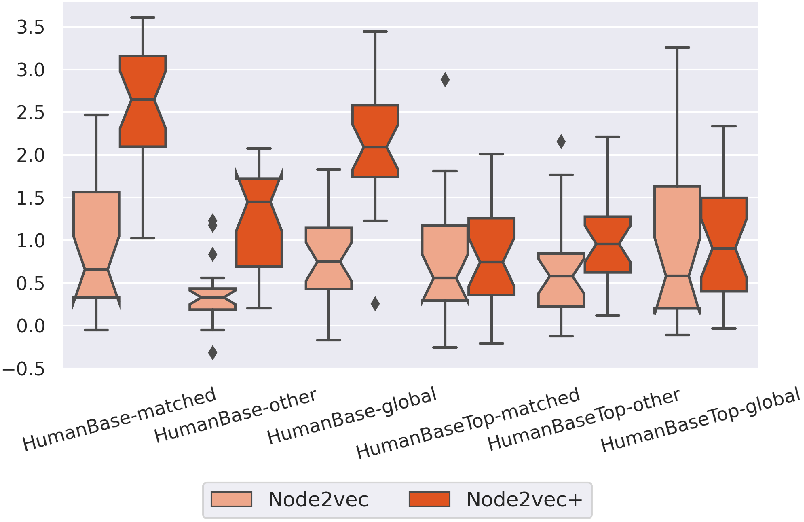
Tissue-specific functional gene classification tasks results.

We observe similar results using another collection of tissue-specific co-expression networks, **GTExCoExp**, that are generated using a benchmarked co-expression network generation workflow by Johnson and Krishnan (2022) (see supplement).

## 4 Discussion and conclusion

In this paper, we propose *node2vec+* that improves upon the second-order random walk in *node2vec* for weighted graphs by considering edge weights. Consequently, the corresponding node embeddings are improved whenever optimal in-out walks positively influence the task (meaning that the optimal *q* setting is not 1).

We illustrated the ability of *node2vec+* to identify potential out edges on weighted graphs, as opposed to *node2vec*, using synthetic datasets including the barbell graph, as well as the hierarchical cluster graphs. Furthermore, evaluations on a variety of challenging gene classification tasks demonstrated that embedding methods like *node2vec(+)* are superior to GNNs. GNNs learn how to orient the nodes in a low dimensional space to maximize the separation between nodes of different classes in an end-to-end fashion. The suboptimal GNN performance here highlights their need for a much larger labeled training dataset to fully exploit the expressive power of their architectures. Unfortunately, many real world biological applications, such as the function or disease gene classification problems here, still lack large amounts of labeled data. For these applications, an unsupervised approach like *node2vec(+)* may be more suitable as it arranges the latent space purely based on the underlying graph structure, after which a less data-hungry model, such as logistic regression, can be applied to perform the classifications.

Dense weighted graphs are common in biology, directly based on the experiment (e.g., genetic interactions (Costanzo *et al*., 2016)), by construction (e.g., co-expression (Zhang and Horvath, 2005)), or by integrating multiple network datasets sources (Greene *et al*., 2015; Szklarczyk *et al*., 2021). Network embedding has recently found applications in studying co-expression networks, for example, in the context of evolutionary and cross-species network alignment (Ovens *et al*., 2021a,b), cancer prognostic gene identification (Choi *et al*., 2018), and gene functional interaction prediction (Du *et al*., 2019). These applications, especially the ones that leverage dense weighted graphs, are likely to benefit from using *node2vec+*.

Sparsification using a hard threshold is a common technique for dealing with fully-connected weighted graphs like co-expression (Zhang and Horvath, 2005; Du *et al*., 2019). However, finding the optimal cut threshold could be quite challenging (usually relying on heuristics (Ovens *et al*., 2021a)) and such thresholding may change the graph significantly in terms of its spectrum (Spielman and Teng, 2010). *Node2vec+*, on the other hand, provides a flexible alternative to hard thresholding due to its incorporation of the noisy edge threshold parameter *γ*, enabling soft thresholding.

Overall, *node2vec+* is a natural extension of *node2vec* for weighted graphs and has several desirable properties. With its general procedure for biased random walks, *Node2vec+* can be easily adapted into other methods such as KG2Vec (Wang *et al*., 2021b) and Het-Node2vec (Valentini *et al*., 2021) that are built on top of *node2vec. Node2vec+* is available as an open-source software as part of the PecanPy package: https://github.com/krishnanlab/PecanPy.

