## Supplemental Information for "Accurately modeling biased random walks on weighted networks using *node2vec+*"

#### Hierarchical cluster graphs construction details

The hierarchical cluster graphs are constructed by first sampling in a representation space constructed based on the corresponding tree structure and then applying an RBF kernel.

##### Tree construction

We first construct the cluster centroids using a tree structure. A *perfect binary tree* is a binary tree where all nodes except for the leaf nodes have two children, and all leaf nodes have the same level. This definition can be generalized to *perfect K-trees*, in which all the interior nodes have K number of nodes, for K greater than or equal to one. We denote $T_{K,L}$ as the *perfect K-tree* with maximum level $L$. Figure S1 shows the example of $T_{2,2}$, a *perfect binary tree* (or *perfect 2-tree*) with a maximum level of two.

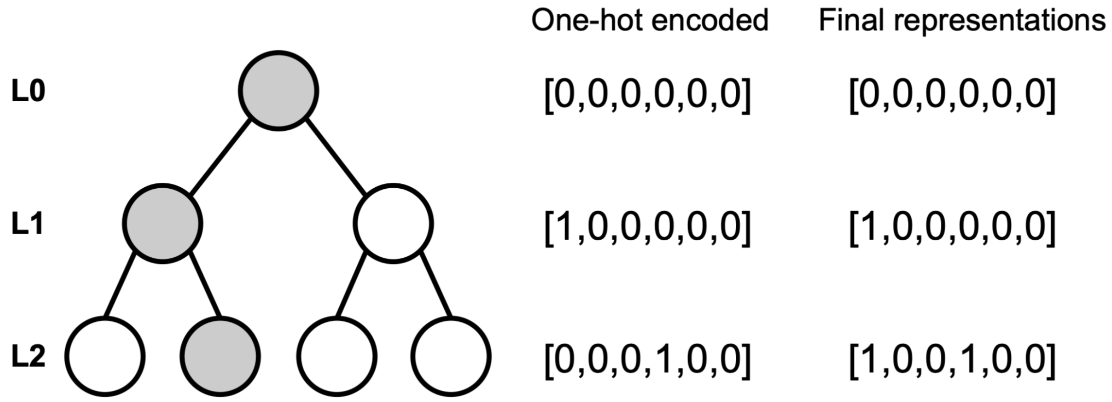


**Figure S1. A perfect binary tree with two levels.**

##### Representing nodes in the tree

A straightforward solution to represent the nodes in $T_{K,L}$ is one-hot encoding. For a more compact representation, we leave out the indicator for the root node, and represent it as all zeros. Thus, the dimension of the indicator array is equal to the total number of nodes in $T_{K,L}$, excluding the root node, which equals to the following

$$|V(T_{K,L})| - 1 = \sum_{l=1}^{L} K^l$$

However, if we use one-hot encoding, then all nodes are equally distanced in the Euclidean space. Instead, we combine the one-hot encoded representations of all the ancestor nodes as the final representation for each node, denoted as $\mu_i$, $i = 1,\ldots, |V(T_{K,L})|$. In this way, all sibling nodes are equally distanced, with $\sqrt{2}$ times the distance from the parent node. Figure S1 shows the example of both the one-hot encoded and the final representations of the grey nodes. Notice the difference between the final representation and the on-hot encoded representation of the grey leaf node.

#### Hierarchical clusters

We draw data points $x$ from a Gaussian distribution around each node in the $T_{K,L}$ tree:

$$x \sim N(\mu_i, \sigma),\ i = 1,..., |V(T_{K,L})|$$

In the case of K3L2, the data points are drawn using $T_{3,2}$. The parameter $\sigma$, which controls the noisiness of the sampled data points, is set to 0.01 by default. Throughout the study, we fix the number of data points per node in the tree to 30. Finally, we turn the sampled data points into a fully connected weighted graph using the RBF kernel.

#### Maximal sparsification of K3L2

We apply a global edge threshold to K3L2 by removing all edges below a certain value. Sweeping through [0.01, 0.9], we found that the maximum global edge threshold that preserves the connectivity of the graph is about 0.45 (Figure S2). Notice that by doing so, the edge density drastically reduces to around 0.1.

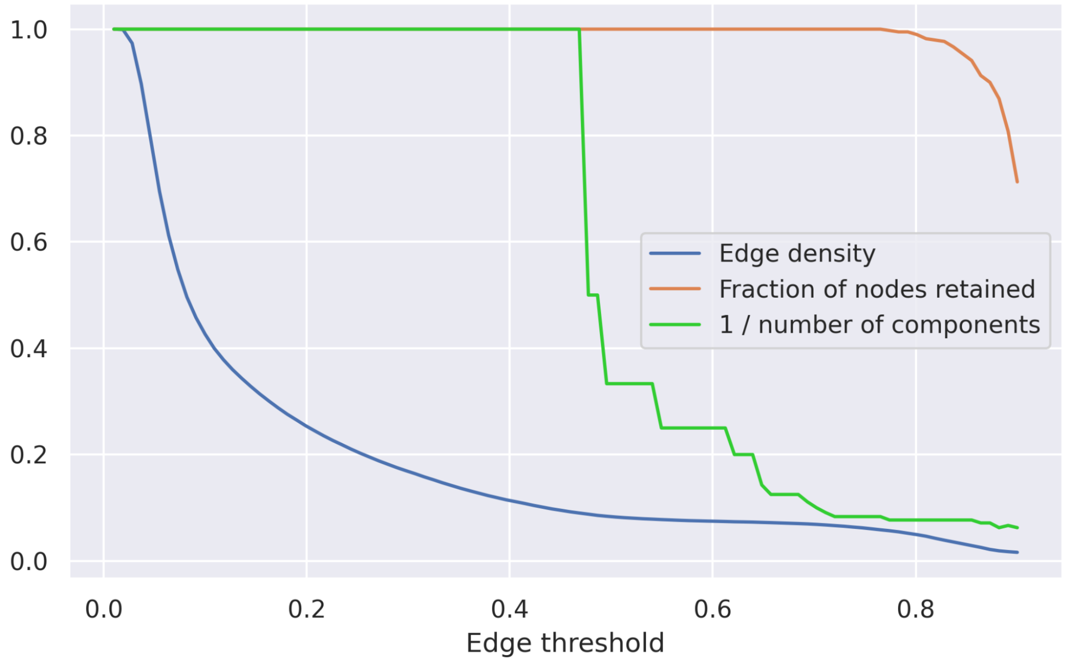


**Figure S2. Network statistics of K3L2 as a function of sparsifying edge threshold.**

### Gene interaction networks statistics

**Table S1. Gene interaction network statistics.**

| Network | # Nodes | # Edges | Edge density |
|---|---|---|---|
| STRING | 17,352 | 3,640,737 | 2.42E-02 |
| HumanBase-global | 24,114 | 290,730,441 | 1.00E+00 |
| HumanBaseTop-global | 24,003 | 33,937,714 | 1.18E-01 |
| HumanBase-blood | 24,114 | 290,730,441 | 1.00E+00 |
| HumanBase-blood_vessel | 24,114 | 290,730,441 | 1.00E+00 |
| HumanBase-brain | 24,114 | 290,730,441 | 1.00E+00 |
| HumanBase-heart | 24,114 | 290,730,441 | 1.00E+00 |
| HumanBase-kidney | 24,114 | 290,730,441 | 1.00E+00 |
| HumanBase-muscle | 24,114 | 290,730,441 | 1.00E+00 |
| HumanBaseTop-blood | 24,003 | 29,451,338 | 1.02E-01 |
| HumanBaseTop-blood_vessel | 24,114 | 55,910,520 | 1.92E-01 |
| HumanBaseTop-brain | 24,003 | 36,199,871 | 1.26E-01 |
| HumanBaseTop-heart | 24,114 | 48,614,273 | 1.67E-01 |
| HumanBaseTop-kidney | 24,114 | 48,599,506 | 1.67E-01 |
| HumanBaseTop-muscle | 24,114 | 54,492,105 | 1.87E-01 |
| GTExCoExp-blood | 19,809 | 95,535,108 | 4.87E-01 |
| GTExCoExp-blood_vessel | 19,809 | 97,085,845 | 4.95E-01 |
| GTExCoExp-brain | 19,809 | 98,573,534 | 5.02E-01 |
| GTExCoExp-global | 20,116 | 98,135,412 | 4.85E-01 |
| GTExCoExp-heart | 19,809 | 97,582,341 | 4.97E-01 |
| GTExCoExp-kidney | 19,809 | 97,929,900 | 4.99E-01 |
| GTExCoExp-muscle | 19,809 | 97,240,974 | 4.96E-01 |
| GTExCoExpTop-blood | 19,809 | 7,884,935 | 4.02E-02 |
| GTExCoExpTop-blood_vessel | 19,809 | 4,879,744 | 2.49E-02 |
| GTExCoExpTop-brain | 19,809 | 6,276,541 | 3.20E-02 |
| GTExCoExpTop-global | 20,116 | 9,959,240 | 4.92E-02 |
| GTExCoExpTop-heart | 19,809 | 5,067,436 | 2.58E-02 |
| GTExCoExpTop-kidney | 19,809 | 8,589,096 | 4.38E-02 |
| GTExCoExpTop-muscle | 19,809 | 5,415,950 | 2.76E-02 |

### GNN training information

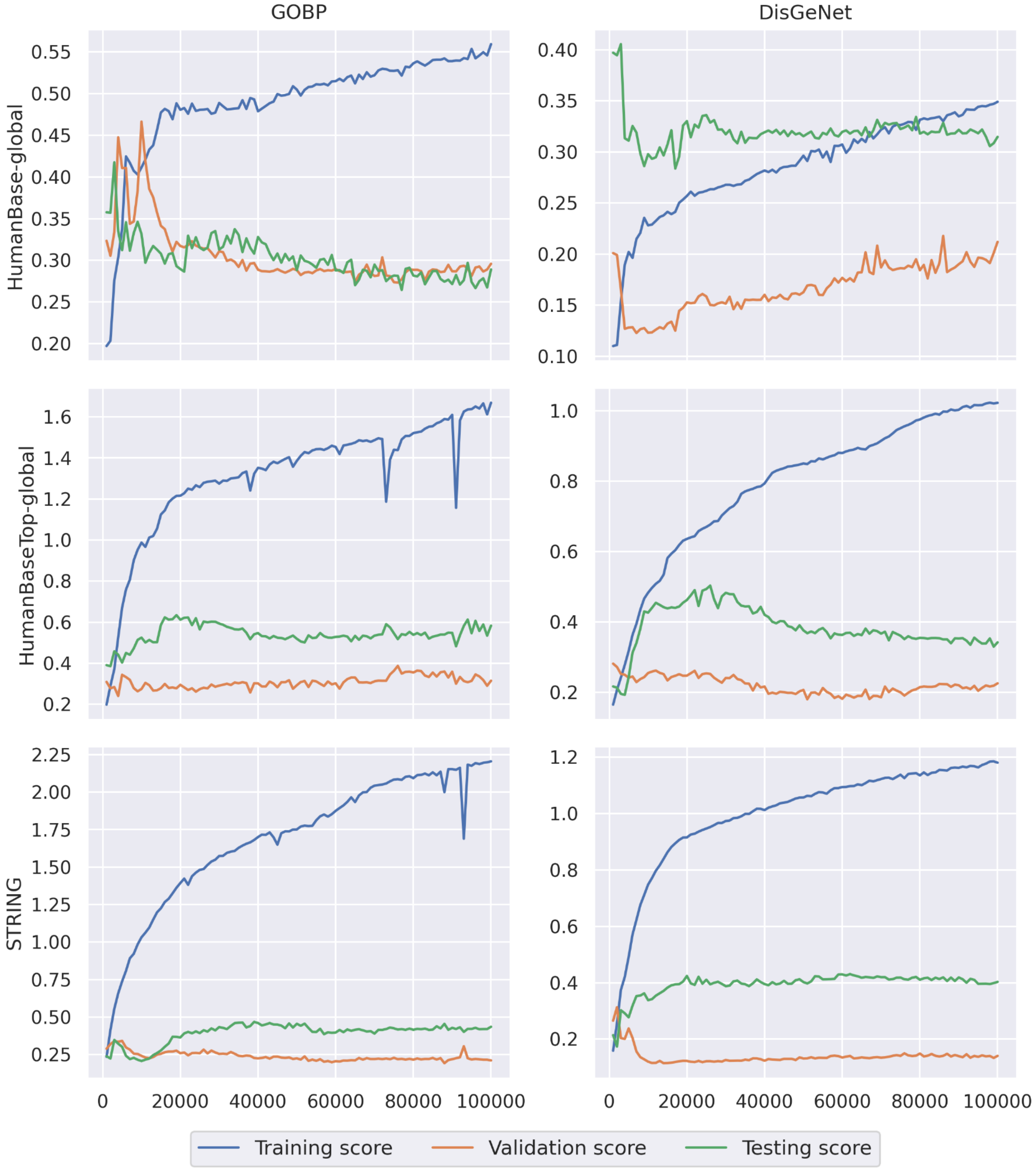


**Figure S3. GCN training progress.**

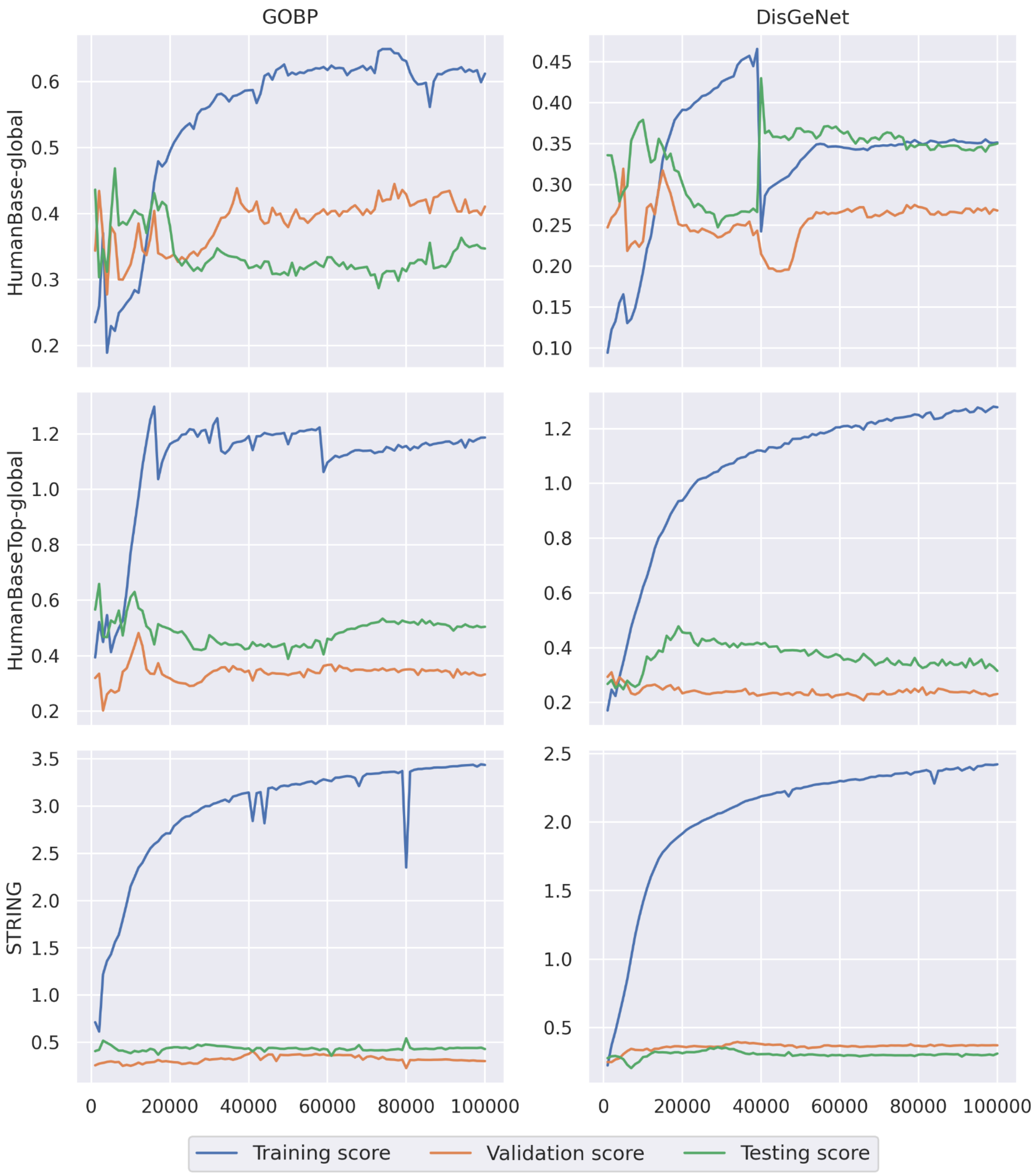


**Figure S4. GraphSAGE training progress.**

### Additional results

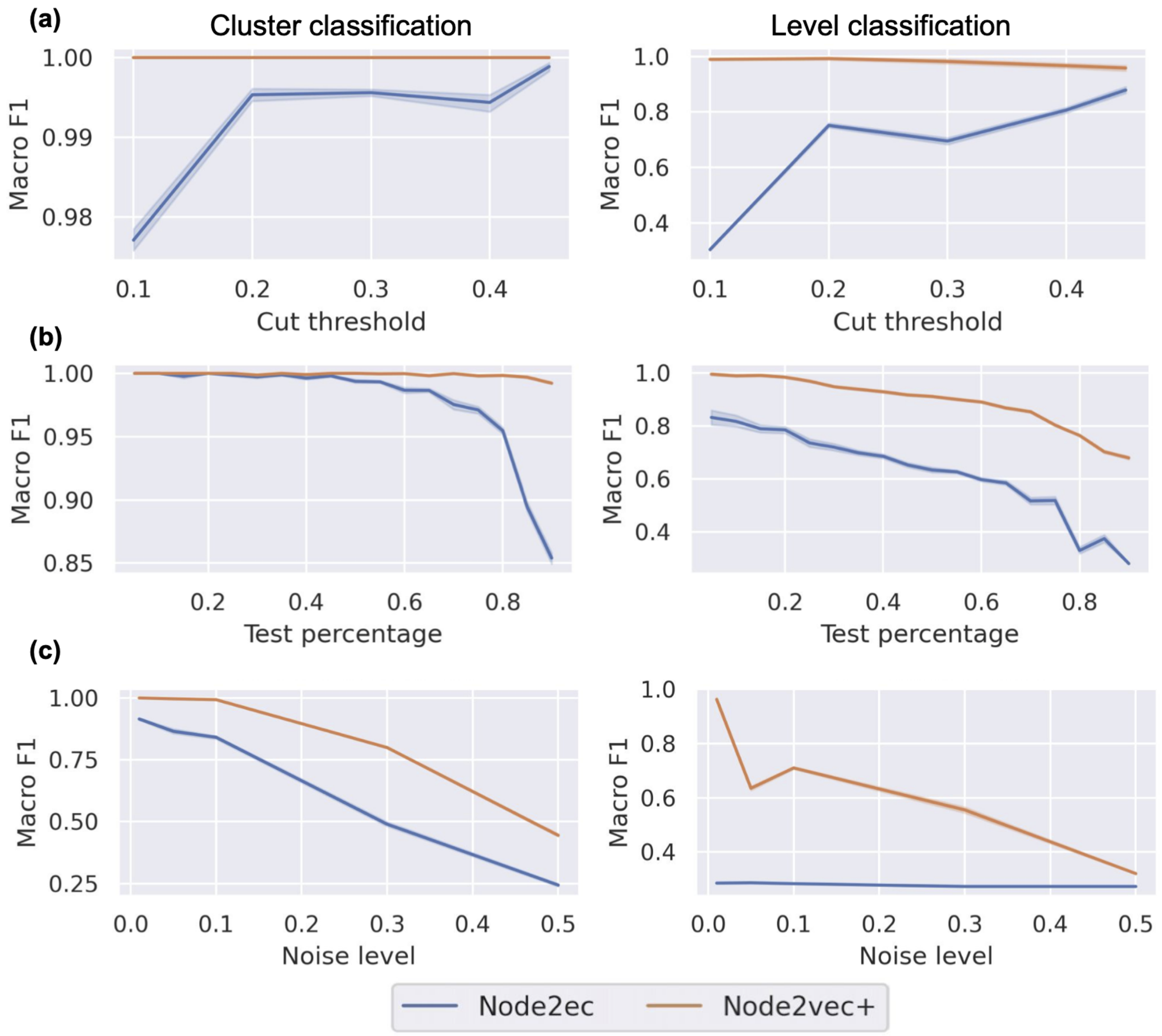


**Figure S5. Fine-grained analysis of K3L2.** (a) changing sparsification threshold value. (b) changing train/test raio, larger value means less training data. (c) changing noise level during network construction.

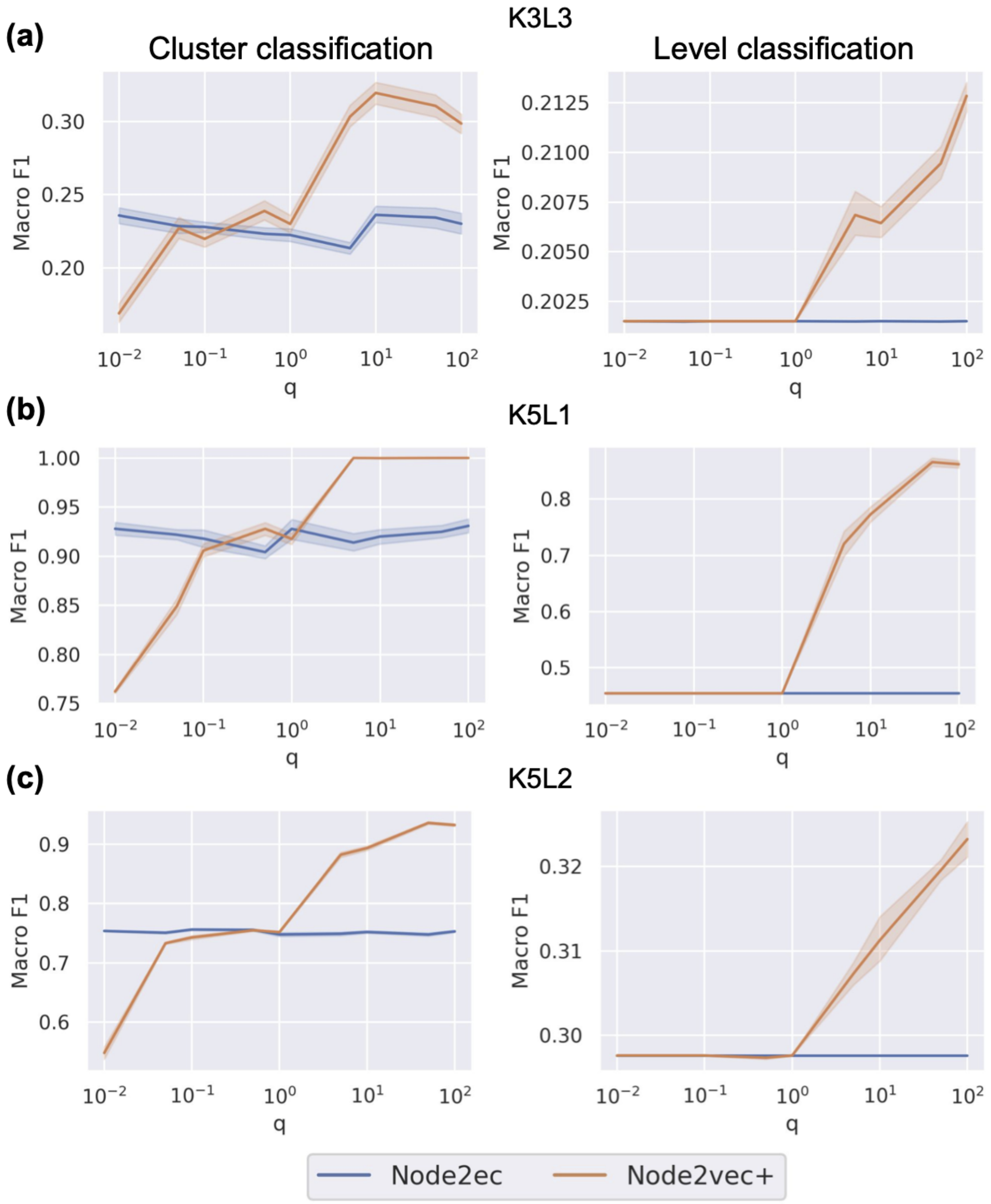


**Figure S6. Evaluation of other hierarchical cluster graphs.**

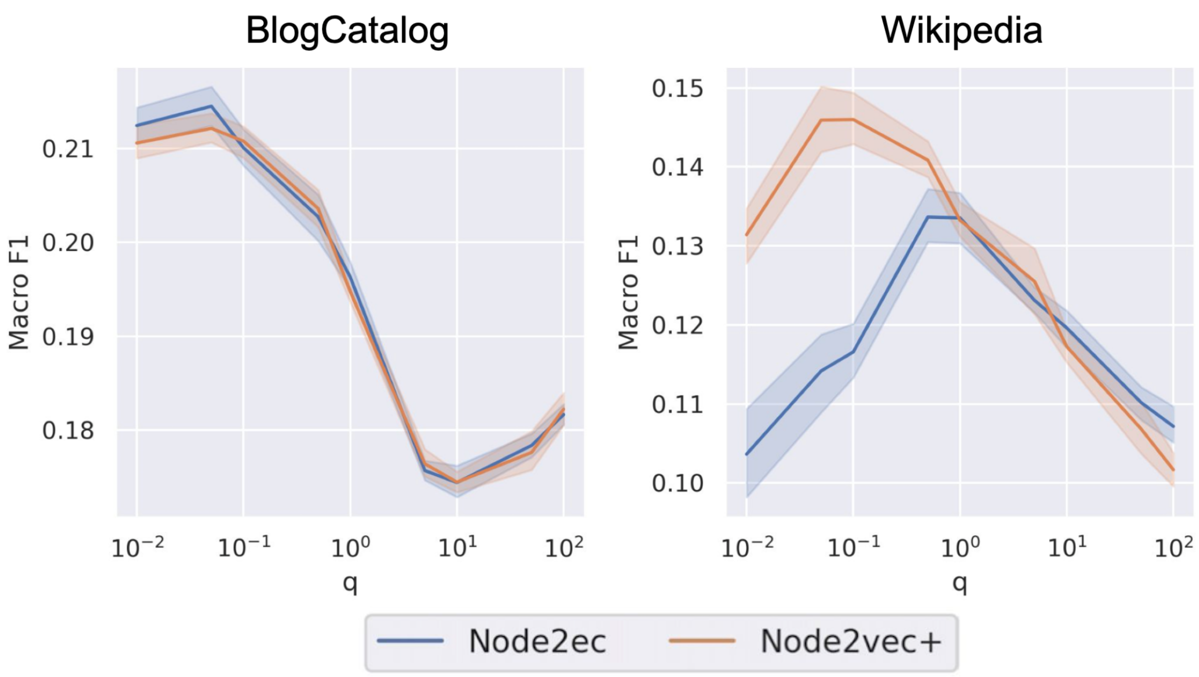


**Figure S7. Multi-label classification benchmarks using BlogCatalog and Wikipedia.** Data are obtained from https://snap.stanford.edu/node2vec

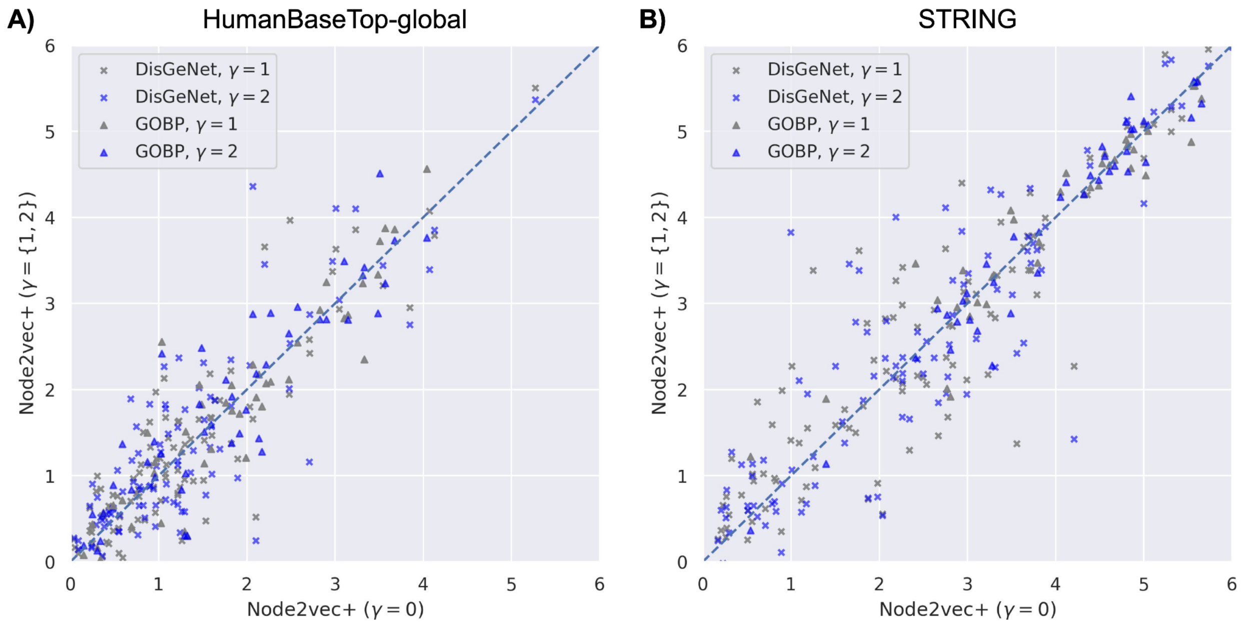


**Figure S8. Effects of gamma on HumanBaseTop-global and STRING**

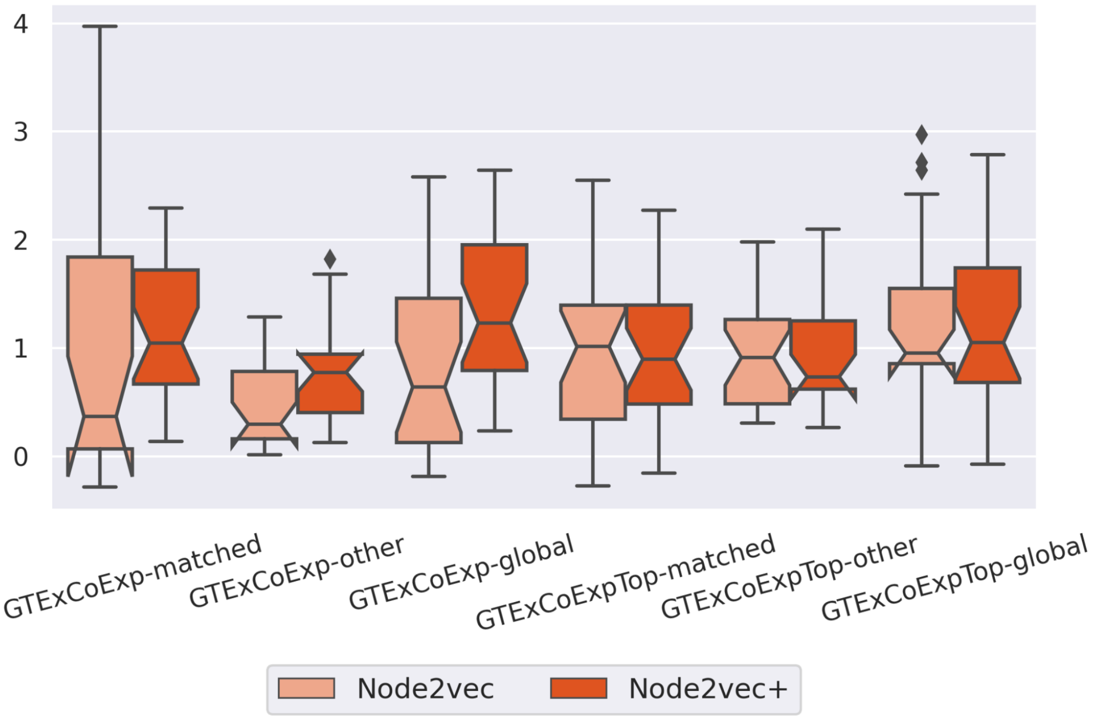


**Figure S9. Tissue-specific functional gene classification tasks using GTExCoExp.**
